# Thymidylate synthase maintains the de-differentiated state of aggressive breast cancers

**DOI:** 10.1101/321943

**Authors:** Aarif Siddiqui, Paradesi Gollavilli, Annemarie Schwab, Maria Eleni Vazakidou, Pelin G Ersan, Mallika Ramakrishnan, Dick Pluim, Si′Ana Coggins, Ozge Saatci, Laura Annaratone, Jan HM Schellens, Baek Kim, Irfan Ahmed Asangani, Suhail Ahmed Kabeer Rasheed, Caterina Marchiò, Ozgur Sahin, Paolo Ceppi

**Author notes:** **Address correspondence to:** Dr. Paolo Ceppi, Junior Research Group 1, Interdisciplinary Center for Clinical Research, Friedrich-Alexander-University (FAU) Erlangen-Nürnberg, Nikolaus-Fiebiger-Zentrum, Glückstrasse 6, 91054 Erlangen, Germany.

## Abstract

Cancer cells frequently boost nucleotide metabolism (NM) to support their increased proliferation, but the consequences of elevated NM on tumor de-differentiation are mostly unexplored. Here, we identified a role for thymidylate synthase (TS), a NM enzyme and established drug target, in cancer cell de-differentiation and investigated its clinical significance in breast cancer (BC). *In vitro*, TS knockdown increased the population of CD24^+^ differentiated cells, and attenuated migration and sphere-formation. RNA-seq profiling indicated a repression of epithelial-to-mesenchymal transition (EMT) signature genes upon TS knockdown, and TS-deficient cells showed an increased ability to invade and metastasize *in vivo*, consistent with the occurrence of a partial EMT phenotype. Mechanistically, TS enzymatic activity was found essential for the maintenance of the EMT/stem-like state by fueling a DPYD-dependent pyrimidine catabolism. In patient tissues, TS levels were found significantly higher in poorly differentiated and in triple negative BC (TNBC), and strongly correlated with worse prognosis. The present study provides the *rationale* to study in-depth the role of NM at the crossroads of proliferation and differentiation, and depicts new avenues for the design of novel drug combinations for the treatment of BC.

## INTRODUCTION

Tumor de-differentiation contributes to the malignant phenotype of most solid tumors, including breast cancer (BC) (Jogi et al., 2012) (Pece et al., 2010), and in many cases this is achieved through the epithelial-to-mesenchymal transition (EMT) and the cancer stem cell (CSC) programs (Ceppi and Peter, 2014; Magee et al., 2012; Mani et al., 2008; Shibue and Weinberg, 2017; Visvader and Lindeman, 2012). Indeed, EMT and CSC markers are more frequently found in aggressive poorly differentiated (high-grade) tumors in breast (Pece et al., 2010) and other cancers (Barbachano et al., 2010; Ceppi et al., 2010; Jiang et al., 2009; McKeithen et al., 2010; Shi et al., 2013). Understanding the mechanisms controlling the differentiation of BC cells can therefore lead to more effective therapeutics. Nucleotide metabolism (NM) is classically viewed as a motor of cellular proliferation (Lane and Fan, 2015). Cancer cells are, in fact, highly dependent on the *de novo* synthesis of nucleotides to produce sufficient DNA and RNA precursors to support their growth, and some cancer-promoting signaling pathways have been shown to regulate NM (Tong et al., 2009). However, a few recent studies have suggested that nucleotides-generating metabolic pathways may also serve as regulators of cancer stemness (Bageritz et al., 2014; Morgenroth et al., 2014), opening the possibility that some NM enzymes are implicated in cancer cell de-differentiation. Thymidylate Synthase (TS) is the enzyme that catalyzes the conversion of deoxyuridine monophosphate (dUMP) to thymidine monophosphate (dTMP or thymidylate). Since this reaction provides the sole *de novo* pathway for thymidylate production, TS is essential for DNA synthesis and repair, and its absence blocks proliferation and causes cell death (Costi et al., 2005; Wilson et al., 2008). We previously discovered that TS expression is correlated with the EMT phenotype in the NCI-60 transcriptomic database by using a pan-cancer EMT gene ratio (Vimentin/E-Cadherin, VIM/CDH1) (Siddiqui et al., 2017), but the mechanistic involvement of the TS enzymatic activity on EMT/CSCs has never been shown. Here, we report a novel fundamental role of TS in maintaining the de-differentiated phenotype of BC cells and its differential expression in the BC subtypes, with several potential therapeutic implications.

## RESULTS

### TS is a marker of more aggressive and EMT-driven breast cancer

In order to study the association of TS expression with EMT markers in BC, we employed a VIM/CDH1 ratio to classify the BC cell lines belonging to the CCLE dataset (n = 52) into epithelial, mesenchymal or intermediate phenotypes (**Fig. 1A** and **Supplementary Table 1**. Comparing epithelial (VIM/CDH1<2) with mesenchymal cells (> 2), a significantly higher expression of TYMS mRNA was found in the latter (p<0.005, **Fig. 1B**). Then, in order to test TS gene expression in the spectrum of BC patients, we analyzed 3 independent GEO datasets, and found that TS expression was significantly different among the BC subtypes. Normal-like samples or the well-differentiated tumors (like luminal A) exhibited low TS expression, whereas high TS levels were found in basal-like BC (**Fig. 1C**). BC with a basal-like gene signature are primarily triple-negative (TN), and are frequently enriched for markers of CSCs and EMT markers (Taube et al., 2010).

**Figure 1.**
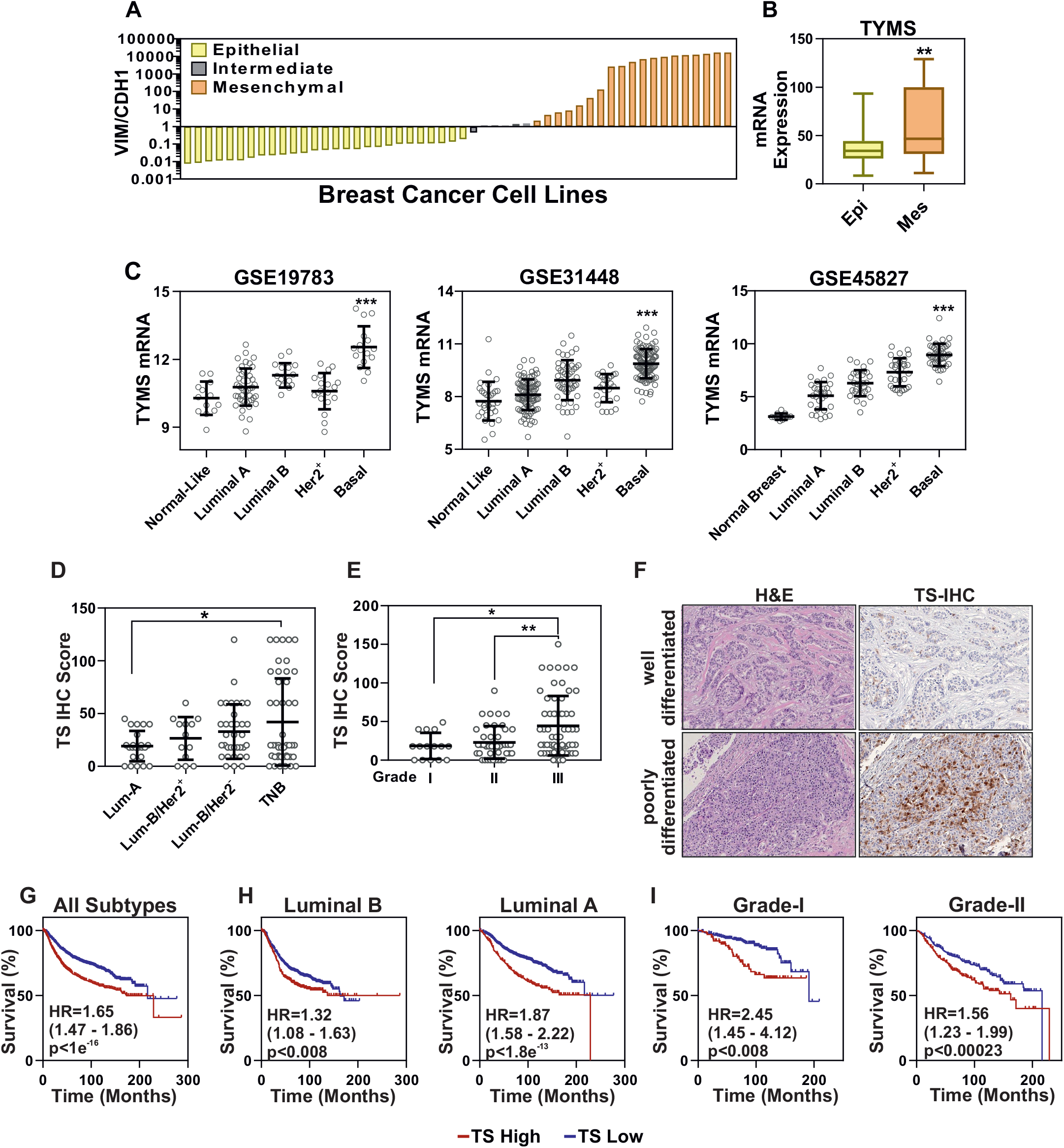
TS is a marker of aggressive and EMT-driven breast cancer. (A) Vimentin/E-cadherin (VIM/CDH1) ration in breast cancer cell lines belonging to the CCLE database (gene expression). Cells were sorted in epithelial (n = 27), mesenchymal (n = 6) and EMT-intermediate (n = 19) phenotype. (B) Comparison of TS mRNA expression levels between epithelial and mesenchymal breast cancer cell lines. P-value is from a two-tailed t test. (C) TS mRNA levels in breast cancer tissues divided by molecular subtypes (data from the indicated datasets). (D) Immunohistochemical staining of TS in FFPE samples from breast cancer patients (n = 120) segregated according to molecular subtype or (E) grade of differentiation (III are poorly differentiated cancers). Y-axis represents TS IHC scores. P-values are one-way ANOVAs. (F) Representative images showing TS staining in well-differentiated and poorly-differentiated tumors. (G) Kapan-Meier curves showing the prognostic significance of TS expression in breast cancer (all types), in luminal (H) and in low grade patients (I). P values are log-rank tests and hazard ratio is represented as HR.

In order to test the clinical significance of the TS protein and its association with the aggressive phenotype in BC, immunohistochemistry (IHC) was performed on formalin-fixed paraffin-embedded samples from 120 BC patients. Patients’ characteristics are shown in the **Supplementary Table 1**. TS staining was quantified by an IHC score and a significantly higher expression was found in the TNBC, as compared to Luminal-A (**Fig. 1D**, ANOVA p<0.01) and in high grade (G3) compared to low grade tumors (**Fig. 1E-F**, ANOVA, p<0.01). A moderate level of correlation was found between TS and Ki67 proliferation marker (Rs = 0.48, **Supplementary Fig 1A**). Analysis of the prognostic values using large patient datasets indicated TS expression as a marker for poor overall survival in BC (all subtypes, **Fig. 1G**). Of note, TS prognostically stratified luminal A and luminal B patients (**Fig. 1H**) as well as lower grade patients (**Fig. 1I**), while no association was not found in more aggressive BC (**Supplementary Fig 1B**).

### TS is essential for the maintenance of a de-differentiated stem-like state in BC

The patient data prompted us to test whether TS plays a role in maintaining the de-differentiated state of TNBC. Hence, we transduced the TNBC cell line MDA-MB-231 with lentiviruses containing non-overlapping shRNA sequences to knockdown TS (**Fig. 2A**). CD44/CD24 surface staining followed by FACS quantification indicated an increase in the population of differentiated CD24^+^ cells, with a concomitant decrease in the CD44^+^ CD24^-^ BC stem cell population (**Fig. 2B**). In order to carefully monitor the effects on cell growth, the confluency of infected cells was examined with real-time proliferation assays upon TS depletion. The results showed a significant suppression of proliferation in cells infected with the shRNA that delivered a strong TS repression (shTS#2), while the sequence with a mild knockdown (shTS#1) did not alter the cells’ growth (**Fig. 2C**). By contrast, a shRNA (shTS#3) sequence that induced a complete TS elimination in this cell line caused a massive growth arrest and death (**Supplementary Fig 1C-D**), in line with the life-essential role of TS. To determine if the partially TS-deficient cells had reduced ability to migrate, we performed a real-time wound-healing assay where cell migration is corrected for proliferation outside the wound to account for changes in cell growth rate upon TS repression. The results showed a significant loss of migratory ability upon TS knockdown (**Fig. 2D-E**). Moreover, assayed for stemness-related functional phenotype, TS-deficient cells formed less mammospheres in low-adherent cultures (**Fig. 2F**). Comparable findings were obtained in other TNBC cell lines, such as BT-549 (**Fig. 2G-J**) and Hs 578T (**Supplementary Fig. 1E-G**), implying that TS could be a common EMT/CSCs regulator in TNBC.

**Figure 2.**
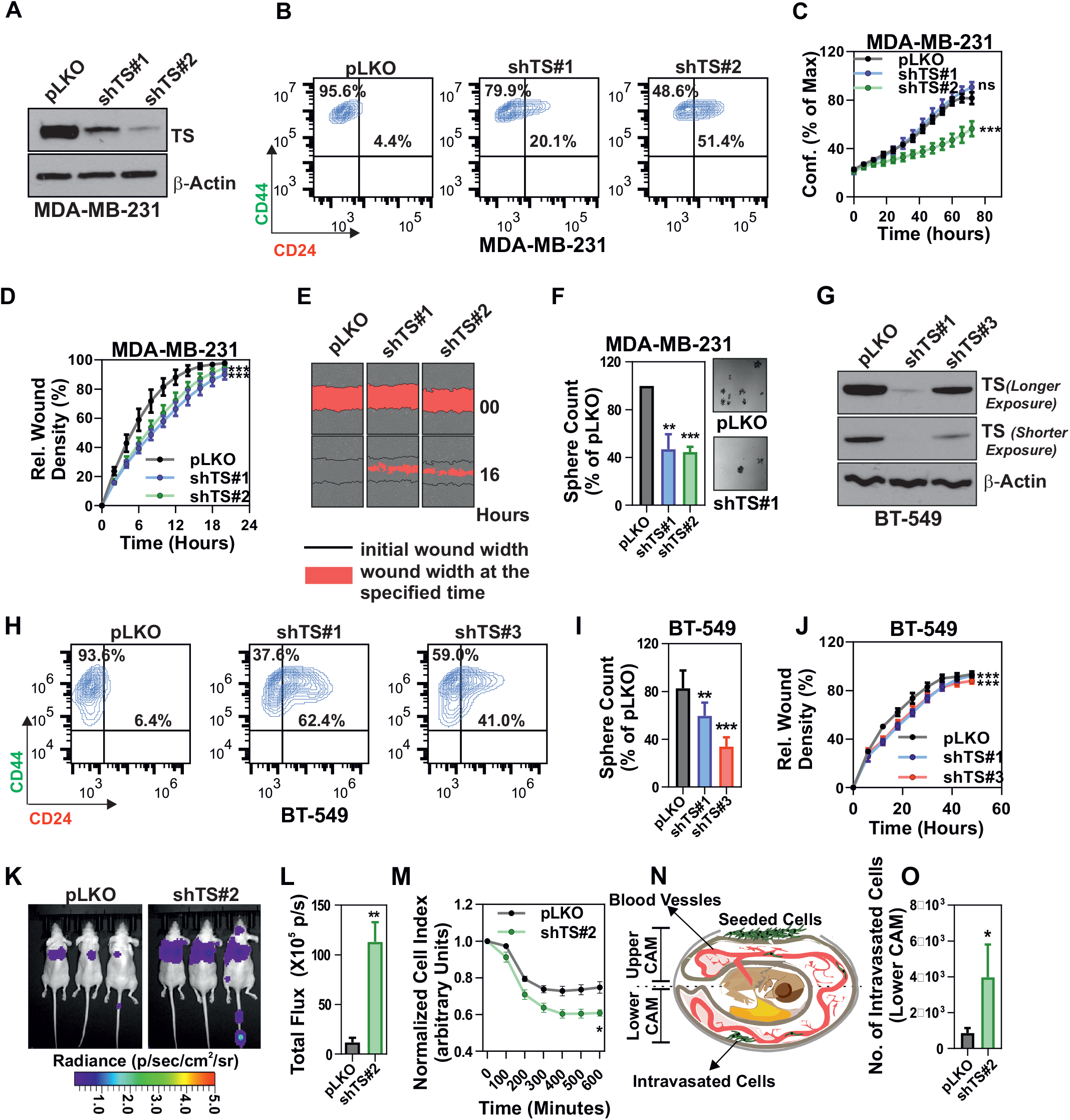
TS knockdown alters the mesenchymal phenotype of TNBCs. (A) Efficiency of TS knockdown in MDA-MB-231 obtained with shRNA lentiviral particles. pLKO is the control non-targeting shRNA. (B) FACS plots of MDA-MB-231 cells as in (A) stained with CD44-FITC and CD24-PE. Gates are based on negative control staining. (C) Real-time measurement of growth (confluency) in MDA-MB-231 with shTS or control cells. (D-E) Real-time migration assay in MDA-MB-231 cells with TS knockdown and (F) relative counts of spheres. (G) Knockdown of TS in BT-549 cells and (H) effect on CD44/CD24 profile. Effect of TS knockdown on (I) sphere-formation capability and (J) migration on BT-549 cells. (K) Representative images of lung metastasis formation assay performed by injecting luciferase-expressing MDA-MB-231 cells in the tail-vein of nude mice (n = 3/group). (L) Quantification of the bioluminescence signal from the lung metastases. (M) Ability of TS knockdown or control cells to penetrate through a HUVEC monolayer (*in vitro* extravasation assay). (N) Schematic diagram representing the CAM assay and (O) quantification of TS knockdown MDA-MB-231 or control cells intravasated in lower CAM 2 days after seeding in the upper CAM. Points are avg±SD. P-values are two-tailed t test (L), multiple t test (M ands O), One-way ANOVA (F, I) or two-way ANOVA (C, D and J). Experimental data are representative from at least two independent experiments with similar results.

Finally, to test the *in vivo* effects of TS suppression, MDA-MB-231-luc cells with TS knockdown or non-targeting control cells were injected in the tail-vein of nude mice. Interestingly, a significantly higher luciferase signal was recorded from shTS cells compared to control cells (**Fig. 2K-L**). Histological analysis confirmed the presence of single or multiple metastatic deposits of various size (ranging from < 1 mm to 2 mm) and signs of lympho-vascular invasion only in mice injected with shTS cells (not shown). The increased ability of TS-deficient cells to form metastasis was independently confirmed by an *in vitro* real-time extravasation assay, which monitors how fast the cancer cells penetrate a monolayer of human umbilical vein endothelial cells (HUVEC) (**Fig. 2M**). In addition, to monitor the intravasation ability, a CAM assay on chicken embryos was performed (**Fig. 2N**), which confirmed an increased ability of TS-deficient cells seeded on the upper CAM to intravasate into the lower CAM (**Fig. 2O**). These observations clearly pointed at a strong impact of TS on BC differentiation and alteration of cells’ behavior, which we aimed to further molecularly characterize.

### TS knockdown induces loss of EMT and correlates with less aggressive BC

To further delineate the molecular pathways regulated via TS and involved in mediating the de-differentiation program in BC, we subjected shTS#1 MDA-MB-231 cells to RNA-sequencing. By setting a stringent cut-off value of 2-fold to identify differentially-expressed genes (DEGs) compared to non-targeting infected cells (pLKO), we found 73 and 84 genes down and up-regulated, respectively (**Supplementary Table 1**). Pathway analysis (GSEA) revealed that EMT was the most significantly deregulated pathway, followed by TNF-α/NFκ-β signaling (**Fig. 3A**), known to be functionally connected with EMT in BC (Li et al., 2012). RNA-seq data were validated by qPCR and (**Fig. 3B, Supplementary Fig. 2A**) were used to explore the prognostic significance of the gene signature associated with TS-knockdown. A system biology approach was adopted to investigate different BC gene expression datasets, dividing the patients by low or high KD score (corresponding to high or low TS, respectively). As a further confirmation of a strong association with EMT, a previously established signature of genes up-regulated during EMT (Sarrio et al., 2008) was found significantly enriched in patients with a low TS KD score (**Fig. 3C**). Moreover, a gene set predicting poor prognosis was found enriched in patients with low KD score (**Fig. 3D**), and similar associations with the grade of differentiation were found using both the TS KD score and TS expression to segregate the patients (**Supplementary Fig. 2B-C**). Survival analysis confirmed the strong prognostic impact of the TS signature, indicating a poorer prognosis for the patients with a lower score or higher TS in all subtypes (**Fig 3E**), and a trend for a significant impact in TNBC (**Fig 3F**). The KD score was also significantly correlated with all the major histopathological variables in BC, including histology, grade of differentiation and size of the tumors (**Fig. 3G-J**). All these data together indicate that TS promotes EMT-driven aggressive BCs.

**Figure 3.**
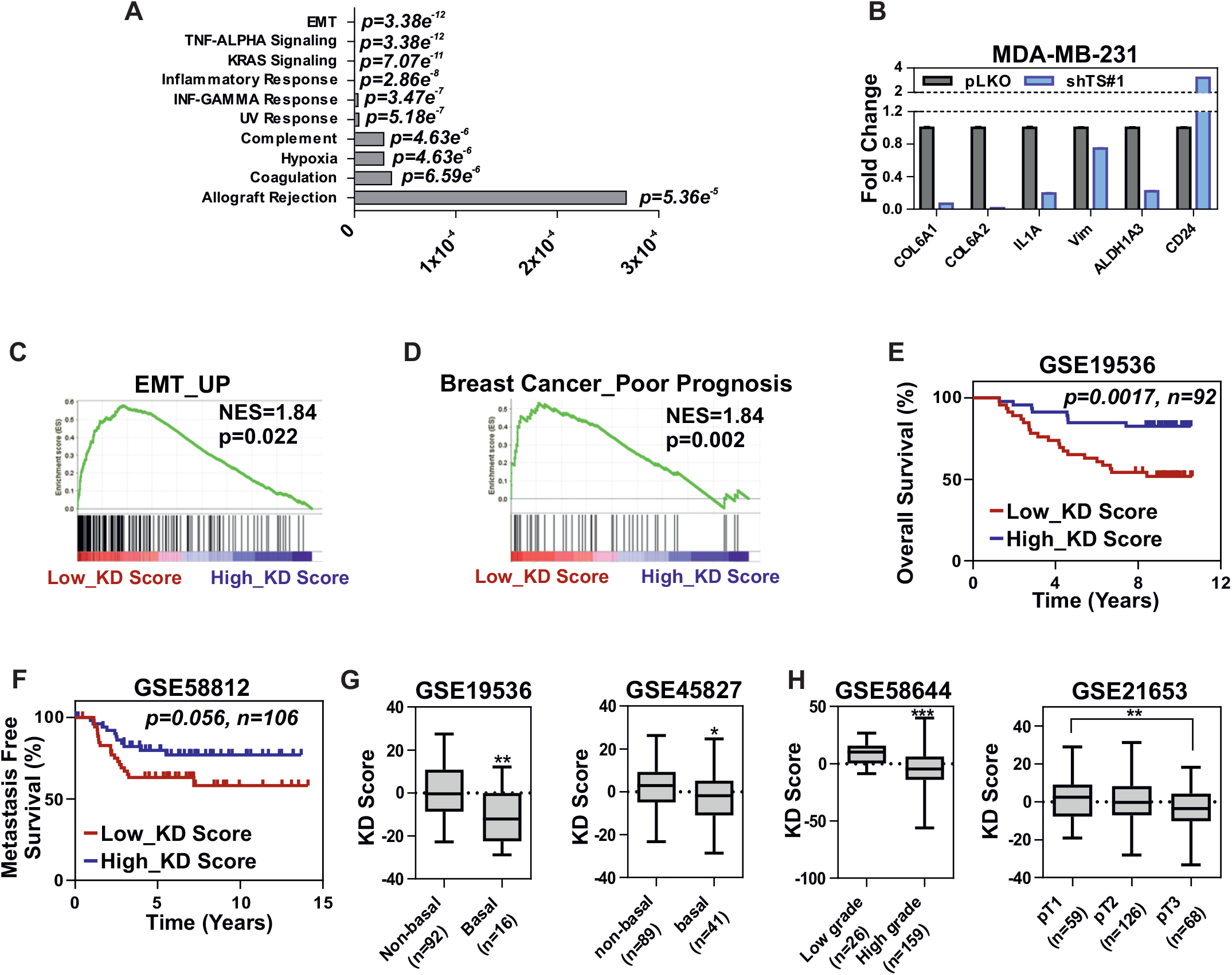
TS knockdown suppresses EMT and the associated gene signature correlates with less aggressive breast cancer. (A) Gene set enrichment (GSEA) analysis showing the most deregulated pathways in MDA-MB-231 with TS knockdown (RNAseq data). (B) qPCR validation of RNA-seq results in MDA-MB-231 shTS#1. Points are avg±SEM. GSEA of TS knockdown scores (low or high TS KD score) with previously published (C) EMT and (D) breast cancer poor prognosis gene signatures, representing NSE (Normalized Enrichment Score) and p –value. Kaplan-Meier curves of (E) overall survival and (F) metastasis-free survival in cancer patients stratified by TS KD score. P values are log-rank tests. TS KD scores in breast cancer patients divided by (G-I) molecular subtype, (I) grade of differentiation, and (J) TNM tumor size. Data are derived from the indicated datasets. P values are two-tailed t test. Experimental data are representative from at least two independent experiments with similar results.

### TS enzymatic activity and thymidine catabolism are essential for maintaining BC de-differentiation

We then aimed at determining the impact of TS enzymatic activity on de-differentiation. TS is the only *de novo* source of thymidylate (dTMP), which is either further phosphorylated to maintain the dNTP pools for DNA synthesis, or directed to degradation via sequential phosphorolytic cleavages. MDA-MB-231 shTS cells with the lowest level of TS knockdown and no major proliferation defect, but with a discernable change in the CD24^+^ population (shTS#1), where subjected to dNTP quantification. Consistent with the previously observed normal growth, no significant change in dTTP or other dNTPs was detected (**Fig. 4A**). This suggested that the differentiation observed in shTS cells was not caused by dNTP imbalance, as shown in other CSC models upon knockdown of specific NM-related enzymes (Bageritz et al., 2014), and that while a baseline TS level is required for the cells to grow, increased TS activity could be partly involved in other cellular functions. To functionally investigate the role of TS enzymatic activity, we reconstituted shTS#1 cells with either a wild type (TS^wt^) expressing silent mutation in shRNA binding region or an enzymatically inactive (TS^R50C^) form of TS (Rahman et al., 2004). Western blotting confirmed the successful transduction (**Fig. 4B**) and TS enzymatic activity assay confirmed the loss of catalytic activity in the TS^R50C^ overexpressing cells (**Fig. 4C**). Strikingly, MDA-MB-231 cells overexpressing the TS^wt^ enzyme increased proliferation (**Fig. 4D**), reduced the population of differentiated CD24^+^ cells (**Fig. 4E**), formed more mammospheres (**Fig. 4F**) and were significantly more migratory (**Fig. 4G**) compared to TS^R50C^ cells overexpressing the catalytically inactive TS. These results clearly indicated that the TS enzymatic activity is essential for the maintenance of the EMT/CSCs phenotype. We therefore hypothesized that TS mediates de-differentiation via the nucleotide catabolism (**Fig. 4H**), rather than through the pathway leading to DNA synthesis. This was supported by the results of a recent shRNA screen on human immortalized breast cells, which revealed that pyrimidine catabolism mediated by dihydropyrimidine dehydrogenase (DPYD) was responsible for the EMT phenotype (Shaul et al., 2014). We therefore speculated the existence of a TS-DPYD axis controlling EMT/CSCs in BC cells. Analysis of expression data from the CCLE dataset reinforced this hypothesis, by showing a marked increase in mesenchymal-like BC cells not only for DPYD, but also for the NT5E (CD73), which is the upstream 5’-nucleotidase responsible for catalyzing the first step of dTMP degradation (**Fig. 4I**). In order to functionally prove our hypothesis, we knocked down DPYD (**Fig. 4J**) and observed a significant increase in the population of CD24^+^ cells (**Fig. 4K**) and a loss of migratory ability (**Fig. 4L**), in line with what we observed in TS-deficient cells. However, the CD24^+^ enriching effect of DPYD knockdown could not be reverted by overexpressing TS (**Fig. 4M**), indicating that a functional DPYD is required for TS to promote de-differentiation. In summary, TS enzymatic activity and the rate of pyrimidine catabolism are essential for the maintenance of the BCSC phenotype. We therefore propose a model in which dTMP produced by TS-overexpressing cancer cells is not only metabolized to support the uncontrolled proliferation, but can also partially sustain de-differentiation and EMT via DPYD-based pyrimidines degradation (**Fig. 4N**).

**Figure 4.**
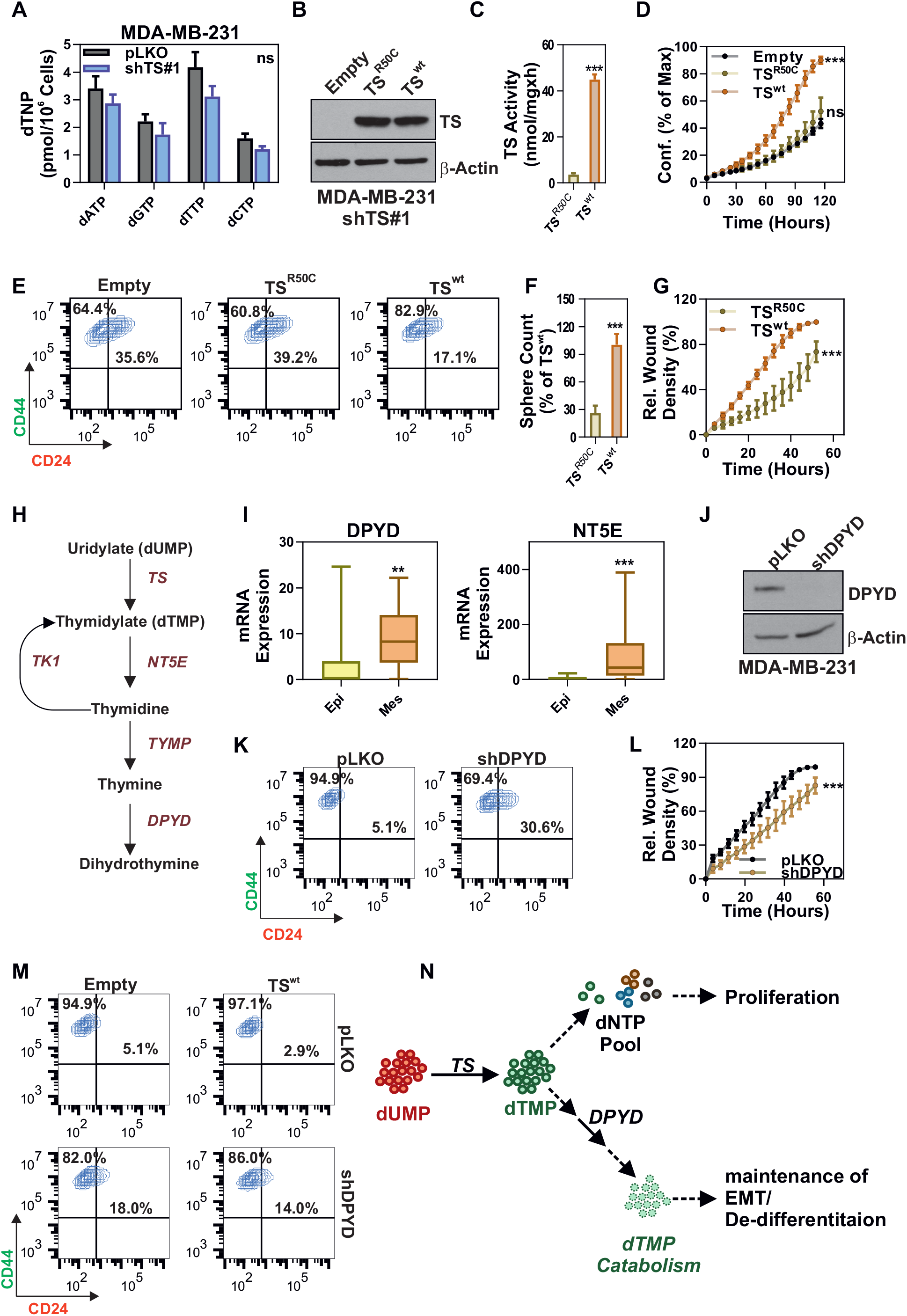
TS enzymatic activity and dTMP catabolism are essential for maintenance of breast cancer de-differentiation. (A) Quantification of deoxyribonucleotide triphosphate (dNTPs) in MDA-MB-231 cells with TS knockdown or control cells. (B) Reconstitution of enzymatically inactive (TSR^50C^) or wild-type (TS^wt^) TS in MDA-MB-231 shTS#1 cells. (C) Measurement of TS enzymatic activity in MDA-MB-231 shTS#1 cells expressing TSR^50C^ or TS^wt^. (D) Proliferation, (E) CD44/CD24 FACS profile, (F) sphere-forming ability, and (G) migration capacity of cells as in (B). (H) Scheme of the thymidylate catabolic pathway. (I) DPYD and NT5E mRNA expression levels in epithelial and mesenchymal breast cancer cell lines (CCLE dataset). (J) Efficacy of DPYD knockdown in MDA-MB-231, and the effects on (K) CD44/CD24 profile and (L) migratory ability. (M) FACS plots of MDA-MB-231 cells with DPYD knockdown overexpressing TS or an empty vector and stained with CD44/CD24. (N) Scheme representing the proposed role of TS in maintaining BC de-differentiation. Points are avg±SD. P-values are two-tailed t test (A, C, F and I) or two-way ANOVA (D, G and L). Experimental data are representative from at least two independent experiments with similar results.

## DISCUSSION

The aggressiveness and the de-differentiated phenotype of neoplastic cells have been strongly connected with alterations in specific metabolic pathways, especially those involved in the transformation of glucose (Colvin et al., 2016; Morita et al., 2018; Schwab et al., 2018; Sciacovelli et al., 2016), while the contribution of other pathways is still majorly unexplored. Elevation in NM is typically associated with the tumor cells’ increased demand for DNA precursors to sustain the uncontrolled proliferation (Lane and Fan, 2015). However, a few studies have shown that some NM enzymes are functionally involved in de-differentiation processes, like the ectonucleotidase ENPP1 in the maintenance of the CSCs-like state in glioblastoma (Bageritz et al., 2014). Similarly, treatment with non-toxic doses of TS-inhibiting drugs was found sufficient to induce a differentiation in multiple myeloma CSCs and sensitize the cells for radiotherapy (Morgenroth et al., 2014). These observations, together with the first demonstration of a connection between TS and EMT (Siddiqui et al., 2017), prompted us to investigate the EMT-driven TNBC model (Mani et al., 2008; Taube et al., 2010). Functional experiments clearly indicates that loss of TS altered the de-differentiated phenotype of TNBCs, reducing CD44^+^ CD24^-^ cells, and suppressing migratory and sphere-forming ability. Consistently, this was accompanied by a robust suppression of EMT-associated genes, as evaluated by RNA-seq analysis. *In vivo*, two independent animal models, confirmed by an *in vitro* extravasation assay, revealed an augmented propensity for intravasation and metastasis formation in cells with TS knockdown. This is in line with previous studies which reported similar results in mesenchymal-like TNBC upon EMT suppression (Le et al., 2014), and a plausible explanation came from studies which found that the tumor-initiating potential might be maximum when TNBC are in a partial EMT (hybrid) state (Bae et al., 2015; Grosse-Wilde et al., 2015; Jolly et al., 2014; Pastushenko et al., 2018; Yu et al., 2013). Our *in vivo* data therefore reflect the complicated nature of EMT, which may also vary in a context-dependent fashion (Celia-Terrassa et al., 2012; Jolly et al., 2017). Targeting TS in other cells, in fact, exerted a clear anti-metastatic effect (Kang et al., 2018). Nevertheless, both the immunohistochemical quantification of TS protein levels and the TS knockdown scores derived from the RNA-seq data were strongly associated with BC de-differentiation and prognosis, indicating a pivotal role of TS in the malignancy of BC. Interestingly, only a moderate co-expression between TS and the proliferation marker Ki67 was found, in line with previous data from other cancers (Ceppi et al., 2008; Monica et al., 2009), supporting our model that TS activity could be implicated in other cellular functions independent of proliferation. We found, in fact, that the EMT/CSC suppression imposed by a level of TS knockdown which did not perturb cells’ proliferation and dNTP balance was rescued by overexpressing wild-type TS, but not by a catalytically inactive mutant. This was particularly important because it ruled out the possible contribution of non-enzymatic activities of TS (Liu et al., 2002), directly pointing at the enzymatic activity as the EMT/CSC driving force. Since dihydropyrimidine accumulation was previously shown to control EMT in BC (Shaul et al., 2014), we tested and demonstrated the hypothesis that high TS enzymatic activity in cancer cells sustains de-differentiation and EMT via a DPYD-dependent pyrimidine catabolism. Several follow-up studies are needed, for instance, to 1) address the impact of additional dTMP-transforming enzymes (like NT5E/CD73) and of salvage pathways; 2) identify the regulatory pathways upstream of the TS-DPYD axis, and the involvement of EMT transcription factors; 3) test a similar function for TS in other malignancies; 4) decipher the contribution of TS dysregulation to cancer-relevant pathways other than EMT (like inflammation). In any case, our data assert that the classical role of TS as a mere proliferation marker needs to be revisited. For instance the previously identified oncogenic and tumor-initiating role of TS (Bertino and Banerjee, 2004; Rahman et al., 2004) could be explained with the direct/indirect control of de-differentiation and stemness. More efforts will need to be dedicated to clarify how the cancer cells balance proliferation and differentiation, and to what exact extent NM genes are implicated in these processes.

TS is a well-established target of chemotherapy, being inhibited by drugs like 5-fluorouracil (5-FU) or by folate analogues (Wilson et al., 2014), and its overexpression in tumors represents a major mechanism of chemo-resistance. From the translational point of view, our finding that TS levels can be significantly different among BC subtypes contradicts earlier works (Pestalozzi et al., 1997) and may be useful to improve the treatment strategies. In our study, TS was found higher in the aggressive BC and in high-grade tumors, in line with previous observations in non-small cell lung (NSCLC) and in gastro-entero-pancreatic cancers (Ceppi et al., 2008; Ceppi et al., 2006; Monica et al., 2009), all representing tumors in which this information was found clinically very important for predicting the efficacy of anti-TS drugs (Scagliotti et al., 2008; Selvaggi and Scagliotti, 2009). Based on the lessons learned from these tumors, the present study and some pivotal works (Lee et al., 2011) can anticipate subtype-specific susceptibilities to anti-TS drugs in BC. For instance, these results may provide the *rationale* for excluding TS-high TNBC patients from anti-TS therapies (Masuda et al., 2017). In addition, the prognostic stratification offered by TS expression may help guide treatment decision making of luminal A and G1/G2 breast cancer patients. Future retrospective and prospective analyses are therefore eagerly warranted.

## MATERIAL AND METHODS

**Cell lines**. TNB cell lines, MDA-MB-231 and BT-549 (Purchased from NCI), were cultured in RMPI-1640 (Sigma) while Hs 578T (purchased from ATCC) were cultured in DMEM (Sigma). Media were supplemented with 10% FBS (Sigma), 1% Pen/Strep (Sigma) and 1% L-Glutamine (Sigma). Cells were profiled for STR, used between passages 3 and 15, examined for the presence of mycoplasma and maintained in Plasmocin (Invivogen) to prevent mycoplasma contamination. Human Umbilical Vein Endothelial Cells (HUVEC) cells were cultured with Ham’s F-12K (Kaighn’s) Medium (Gibco) supplemented with 0.1mg/ml Heparin (Sigma) and 0.03mg/ml Endothelial cell growth supplement (ECGS, Sigma) 50u/ml penicillin/streptomycin (Lonza) and 10% fetal bovine serum (Biowest).

**Patients and samples**. Formalin-fixed paraffin embedded samples of a series of 120 consecutive breast carcinomas collected between 2010 and 2013 at the Azienda Ospedaliera Universitaria Città della Salute e della Scienza di Torino (Turin, Italy) were analyzed. All cases were reviewed and anonymized using an alpha-numerical code. The main patients’ characteristics are shown in **Supplementary Table 1**. The use of retrospective solid tumor tissues for the immunohistochemical study was approved by the Ethic Institutional Review Board (IRB) responsible for “Biobanking and use of human tissues for experimental studies” – Department of Medical Sciences, University of Turin (Italy).

**Immunohistochemistry staining and scoring**. 3μm thick sections of formalin fixed paraffin embedded samples were stained for immunohistochemistry with the EPR4545 antibody (Abcam, 1:150 dilution). Immunohistochemistry was performed using an automated slide-processing platform (Ventana BenchMarck XT Autostainer, Ventana Medical Systems). The antibody was optimized on FFPE cell block sections of MDA-MB-231 cells and on human tonsil tissue. Positive and negative controls were included for each immunohistochemical run. Immunoreactivity was assigned based on the proportion of positive tumor cells over total tumor cells (percent positivity) ranging from 0 to 100%. Staining intensity was evaluated as negative, faint, moderate, and intense. If the staining intensity was heterogeneous, then scoring was based on the greatest degree of intensity. Data were integrated in the H-score, calculated as follows: H-score = ΣPi (i + 1), where i represents the intensity of staining (0–3 +), and Pi stands for the percentage of stained tumor cells (0% to 100%).

**Lentiviral Transduction**. Plasmids for TS knock-down lentiviral particles (TRCN000045663, TRCN000045666 and TRCN000045667) and DPYD knock-down viral particles (TRCN000025799) were purchased from Sigma. Empty vector backbone, pLKO, was used as knock-down control. For TS reconstitution, plasmids were purchased from GeneCopoeia in which silent mutation was introduced in shTS#1 binding region (5’-G_483_CAAAGAGTAATCGATACAAT_503_-3’) of TYMS (NM_001071). Enzymatically inactive TS expression vector was generated by introducing single point mutation (5’-C_148_GC_150_-3’ → 5’-TGC-3’) in reconstitution vector. Empty backbone vector was used as control.

For production of lentiviral particles 293T cells were transfected with 8μg knock-down/expression vectors and 2μg of each packaging vectors (pMDL, pV_s_V_g_ and pRevRes) in complex with 24μg PEI (Polysciences). Viral titer was allowed to concentrate for 48 hours, after which supernatant was collected, centrifuged and passed through 0.22μm syringe filter. For transduction, cells were seeded in 6-well plate (100,000 to 150,000) and infected with each lentivirus at a multiplicity of infection (MOI) of 3 in the presence of 8μg/ml polybrene (Sigma). After 48 hours of infection, cells were selected in medium containing 3μg/ml puromycin (Sigma) and cultured in 1 μg/ml puromycin. MDA-MB1–231 cells infected with TS reconstitution vectors were selected in 800μg/ml G418 (Sigma) and cultured in 250μg/ml G418.

**Quantitative Real-Time PCR**. Cells were grown to 80–90% confluency, and were lysed in 700μl Qiazol (Qiagen). Total RNA was extracted using miRNeasy kit (Qiagen) following the manufacturer’s instructions. mRNA was converted to cDNA using Tetro cDNA synthesis kit (Bioline) with random hexamers and 50ng cDNA was used as template for real-time quantification. GAPDH was used as the internal control. TaqMan probes (Thermo-Fisher) were used for quantification in Applied Biosystems 7300. SDS software was used to extract raw data and fold change was calculated using the ΔΔCt method.

**RNA Sequencing**. Total RNA was extracted using miRNeasy kit (Qiagen) following the manufacturer’s instructions. RNA-Seq libraries were constructed using the TruSeq sample Prep Kit V2 (Illumina) according to the manufacturer’s instructions. Briefly, 1μg of purified RNA was poly-A selected and fragmented with fragmentation enzyme. After first and second strand synthesis from a template of poly-A selected/fragmented RNA, other procedures from end-repair to PCR amplification were done according to library construction steps. Libraries were purified and validated for appropriate size on a 2100 Bioanalyzer High Sensitivity DNA chip (Agilent Technologies.). The DNA library was quantified using Qubit and normalized to 4nM before pooling. Libraries were pooled in an equimolar fashion and diluted to 10pM. Library pools were clustered and run on Nextseq500 platform with paired-end reads of 75 bases, according to the manufacturer’s recommended protocol (Illumina). Raw reads passing the Illumina RTA quality filter were pre-processed using FASTQC for sequencing base quality control. Sequence reads were mapped to UCSC human genome build using TopHat and differential gene expression determined using Cufflinks 2.1.1 and Cuffdiff2.1.1 as implemented in BaseSpace (https://basespace.illumina.com/home/indexIllumina).

**Western blot analysis**. Cells were lysed in RIPA lysis buffer and quantified using the Pierce BCA protein assay kit (both from Thermo-Fisher). Proteins lysates (10– 35μg) were resolved on 10% SDS–PAGE gels and transferred to PVDF membrane (Thermo-Fisher). Membranes were blocked in 5% Milk (BioRad) in 1XTBS-T. Membranes were then incubated in primary antibodies diluted in blocking solution at 4°C overnight. Western blot antibody for TS (clone: EPR4545) and DPYD (clone: EPR8811) are from Abcam and β-Actin (clone: 8H10D10) is from Cell Signaling. After incubation with secondary antibodies (Southern Biotech), detection was performed using the ECL method (Pierce ECL western blotting substrate, Thermo-Fisher) and developed on X-Ray film (Thermo-Fisher) using a chemiluminescence imager, AGFA CP100.

**Cell Surface Staining and FACS**. Anti-CD44-FITC and anti-CD24-PE antibodies were purchased from Biolegend. For CD44/CD24 staining, 400,000 cells were plated in 6-wells and allowed to grow overnight. Cells were trypsinized and collected in FACS tubes. After two washes with 1% FBS/PBS, cells were incubated with 400ng CD24-PEA and CD44-FITC (diluted in 100μl 1% FBS/PBS) for 20 minutes at ice. Cells were again washed with 1% FBS/PBS twice and suspended in FACS resuspension buffer (5mM EDTA and 2% FBS in PBS). CD24/CD44 positivity was recorded in CytoFLEX flow cytometer. Auto-fluorescence was recorded by reading the respective unstained cells and was used to gate for CD24 and CD44 positive populations. FACS data were analyzed using FlowJo (V10.1).

**Proliferation Assay**. For Proliferation assays cells were seeded in 96 well plates in low density (5–20% initial confluency). Plates were loaded in Incucyte-Zoom (Essen Bioscience) and scanned every 2–4 hours. For each scan, phase contrast image was acquired from each well and was analyzed by Incycuyte Zoom software to calculate the confluency percentage for the given time.

**Mammosphere Culture**: 40,000 cells were seeded in triplicates in ultra-low attachment 6-well plates (Corning) in complete Mammocult medium (Stem Cell Technologies), prepared according to the manufacturer’s instruction. After formation of visible spheres were counted by spinning at 300g for 5 minutes and suspending in PBS (Lonza).

**Migration Assay**. For migration assays cells were plated in 96 well plates so that they reach 100% confluency overnight. Cells were wounded using WoundMaker (Essen Biosciences) as per the instruction from the manufacturer. Plates were loaded in Incucyte Zoom and were automatically scanned for programmed time interval. For each scan, wound width was recorded by the software and was normalized to the proliferation outside the wound, giving relative wound density for each time point.

**Mouse Tail Vein Metastasis Injection**. For tail vein metastasis assay, 1.5 × 10^6^ cells were injected into tail vein of 6–8 weeks old female athymic nu/nu mice, with 3 mice per group. Lung metastases were monitored by bioluminescence imaging (BLI). Anesthetized mice were intraperitoneally injected with 200 mg/kg D-luciferin (Perkin Elmer). Bioluminescence images were acquired with Lumina III *in Vivo* Imaging System (Perkin Elmer). Analysis was performed with live imaging software by measuring photon flux. Lungs were collected and fixed in 10% buffered formalin and processed to obtain paraffin blocks. Three-micron thick sections of formalin fixed paraffin embedded samples were stained using Hematoxylin and eosin (H&E). The mice experiments were performed at the Bilket University Ankara and were approved by Animal Ethics Committee of the university.

**Real Time HUVEC Invasion Assay (*in vitro* Extravasation Assay)**. *in vitro* extravasation assay was performed as previously described (Au – Rahim and Au – Üren, 2011). Briefly, 2.5X10^4^ HUVEC cells in 100μl were seeded in E-plates. Once HUVEC monolayer formed (19–21 hours), 1X 10^4^ MDA-MB-231 cells were added on top of the monolayer. A decrease in cell index indicates invasion through the HUVEC monolayer by tumor cells. Penetration of HUVEC monolayer by MDA-MB-231 cells was monitored for 6–9 hours by using xCelligence Real-time Cell Analyzer (RTCA) system (Acea Biosciences, San Diego, CA, USA).

**Chick Chorioallantoic Membrane (CAM) Assay**. CAM assay for intravasation was performed as described previously (Rasheed et al., 2010). Briefly, Fertilized White Leghorn eggs (Asby) were incubated in a rotary incubator at 37°C with 60% humidity for 10 days. The CAM was dropped by drilling a small hole through the eggshell into the air sac and a second hole near the allantoic vein that penetrates the eggshell membrane but not the CAM. The CAM was dropped by applying a mild vacuum to the hole over the air sac. Subsequently, a cutoff wheel (Dremel) was used to cut a 1 cm^2^ window, encompassing the second hole near the allantoic vein to expose the underlying CAM. The CAM was gently abraded with a sterile cotton swab to provide access to the mesenchyme and 50μl inoculum of 2X10^6^ MDA-MB-231-pLKO (n = 10) or MDA-MB-231-shRNA-TS#2 (n = 10) cells were applied. 4 eggs without addition of any human cells is used as Negative control (n = 4). The windows were subsequently sealed and the eggs were returned to a stationary incubator. The eggs remained in the incubator for further 2 days, following which the CAM (on day 3) was cut into two halves and the tumor was excised from the upper CAM and the genomic DNA was extracted from the lower CAM. The lower CAM genomic DNA was then analyzed for the presence of human tumor cells in the background of chicken DNA by quantitative TaqMan *Alu* PCR with 500ng gDNA. The number of intravasated human cells was then plotted in the graph as shown.

**Deoxynucleotide Triphosphate Quantification**. The cellular dNTP levels were determined by the RT-based dNTP assay (Diamond et al., 2004). Briefly, the cellular dNTPs in experimental triplicates were extracted by methanol, and the determined dNTP amounts were normalized for an equal cell number (1×10^6^).

**TS Enzyme Activity Quantification**. TS was quantified in MDA-MB-231 cells as previously described (Pluim et al., 2013Pluim et al., 2013). Briefly, cells were collected and pelleted. cells were suspended in 300 μl ice-cold Reaction mix (RM, 20mM MgCl_2_, 1.5mM NaF, 1mM DTT in 50mM Tris-HCl pH 7.5, after deoxygenation 0.47 (v/v%) BME was added. Next, cell lysates were prepared on ice by applying 15 pulses with a Branson 250 tip sonicator (Branson) at power input setting level 3 with a 50% duty cycle. After centrifugation at 11000g for 20 min at 4°C, 95μl of supernatant was transferred to a clean 1.5 mlvial on ice for immediate determination of protein followed by TS activity analysis. Protein concentrations in PBMC cytosolic lysates were determined using the Bio-Rad protein assay (Bio-Rad). Briefly, 5μl of PBMC cytosolic lysate was diluted with 45μl of MilliQ water (Millipore). Five bovine serum albumin standards were prepared in concentrations ranging from 32.5 to 500mg/ml to obtain a standard curve. In duplicate 10μl of diluted lysate and the standard curve were transferred to a clear 96-well flat bottom plate. After the addition of 200μl dye solution, the plate was incubated for 15 min at RT and subsequently the absorption was measured at 590nm using an EL340 microplate reader (Bio-Tek). Immediately before the start of TS activity assay, a vial containing 2.51mg of lyophilized MTHF was reconstituted in 500μl of deoxygenized water and 10μl was added to a 2.0ml vial on ice. To this vial 85μl of ice-cold tumor cell cytosolic lysate corresponding to 15μg of protein was added. Next, 5μl of 1mM ice-cold substrate was added, and after mixing, the samples were incubated for 3 hours at 37°C in a shaking water bath. The reaction was terminated by adding 100μl of 6.5N HCl, and the remaining substrate was bound onto 400μl Carbon slurry (CS, 5g acid washed charcoal, 50mg Dextran T500 in phosphate buffered saline) by vertical disk rotation mixing of the samples at 50 rpm at 4°C. After centrifugation at 11,000g for 5 min at 4°C, 300μl of clear supernatant was transferred to a 20ml polyethylene vial, mixed with 10ml of Ultima Gold, and subsequently assayed for radioactivity for 5 min using a LSC2900 Tri-Carb liquid scintillation counter.

**Bioinformatic analysis**. Gene set enrichment analysis (GSEA) on the differentially expressed genes upon TYMS knockdown was carried out with the Molecular Signatures Database v6.1 software (Broad Institute). For patient data analysis normalized gene expression data from the following patient datasets were downloaded from GEO database; GSE19536, GSE58644, GSE21653, GSE45827 and GSE58812. While GSE58812 is a dataset of triple-negative breast cancer patients, all other datasets are of breast cancer patients with different subtypes. For calculation of TYMS Knockdown (KD) score, first, z scores of the down and up regulated genes upon TYMS knockdown were calculated. Then, the sum of z scores of downregulated genes were subtracted from the sum of z scores of upregulated genes and KD scores were obtained for each patient. Patients were grouped based on either their TYMS gene expression or KD score. GSEA was performed by using patient data from GSE58644 and GSE58812. Survival graphs were generated in GraphPad and the significance was assessed by Log-rank test. Survival graphs from the KM Plotter database was generated based on TYMS expression by using the auto select best cutoff option. Statistical analyses were performed by unpaired student’s t-test. ^*^, p<0.05; ^**^, p<0.01.

**Statistical analysis**. Statistical tests were performed with the GraphPad software v.7 comparing groups of different conditions with replicates. In all tests, the statistical significance was set at p = 0.05 (in the figures ^*^ indicates p<0.05, ^**^ indicates p<0.01, ^***^ indicates p<0.001).

## Acknowledgments

Work supported by the Interdisciplinary Center for Clinical Research of the University of Erlangen-Nuremberg (PC), NIH GM104198 and AI136581 (BK), NMRC/BNIG/2041/2015 (SAK), Fondazione Umberto Veronesi (LA), MIUR 2015HAJH8E (CM).

## Authors’ contribution

AS, PG, ASchwab, MEV, PGE, MR, DP, SAC, OSaatci, and LA performed the experiments and analyzed the data. PG and IA performed and analyzed the RNA sequencing experiments. PGE and OS performed mouse experiments. SAKR performed chicken-embryo assays. MR, LA, CM provided human paraffin-embedded materials, performed IHC, histological evaluation and IHC scoring. AS, IA, JHMS, BK, OS, CM, and PC critically discussed the work. AS and PC designed the experiments and wrote the paper.

